# Identification of cortical targets for modulating function supported by the human hippocampal network

**DOI:** 10.1101/2024.06.02.597047

**Authors:** Hsin-Ju Lee, Fa-Hsuan Lin

## Abstract

**Background:** Individualized transcranial magnetic stimulation (TMS) targeting using functional connectivity analysis of functional magnetic resonance imaging (fMRI) has been demonstrated to be advantageous in inducing neuroplasticity. However, how this approach can benefit modulating the episodic memory function supported by the hippocampal network remains elusive.

**Objective:** We use the resting-state fMRI data from a large cohort to reveal tentative TMS targets at cortical regions within the hippocampal network.

**Methods:** Functional MRI from 1,133 individuals in the Human Connectome Project was used to analyze the hippocampal network using seed-based functional connectivity.

**Results:** Using a weighted sum of time series at the cortex, we identified the average centroids of individualized targets at the medial prefrontal cortex (mPFC) and posterior parietal cortices (PPC) at (-10, 49, 7) and (-40, -67, 30) in the left hemisphere, respectively. The mPFC and PPC coordinate at the right hemispheres are (11, 51, 6) and (48, -59, 24) in the right hemisphere, respectively. These coordinates can be reliably identified (>90% of individuals).

**Conclusion:** Our results suggest candidate TMS target coordinates to modulate the hippocampal function.

## Introduction

Memory performance degrades with age [1]. The early cognitive decline in Alzheimer’s disease (AD) is characterized by the inability to form new memories of events and experiences. Both lesion [2] and functional imaging studies [3] have convergently demonstrated a large-scale network within the medial temporal lobe to be the main neural substrate for memory function [4]. In this network, the hippocampus is a hub that plays a critical role in memory encoding, consolidation, and recollection [5]. Transcranial magnetic stimulation (TMS) can non-invasively stimulate neurons in the superficial but not deep cortex or subcortical regions. Deep brain areas can, however, be activated indirectly by TMS delivered to connected cortical areas [6]. Brain regions supporting episodic memory include the hippocampus, parahippocampal gyrus, retrosplenial cortex, the posterior parietal cortex (PPC) and medial prefrontal cortex (mPFC) [7]. TMS to specific frontal and parietal areas will likely activate the hippocampus via their connectivity.

The distribution of the TMS-induced electric field strength is focal [8]. TMS effects alter significantly as stimulating targets separate by a few millimeters. Thus, TMS targeting should be spatially accurate to bring the optimal modulation. Recent studies suggest that tailoring TMS targeting for each individual can improve the efficacy [9]. Specifically, functional magnetic resonance (fMRI) in the resting state has been used to delineate cortical areas with brain dynamics correlated with that of the interested “regional seed” [10]. This approach has been extensively studied to reveal the dorsolateral prefrontal cortex (DLFPC) functionally connected to the sub-genual anterior cingulate cortex to treat major depression disorder [10–12]. Furthermore, TMS targeting the DLPFC based on individual’s fMRI can improve the response with respect to a common target [13].

The episodic memory function can also be modulated by TMS. A recent review reported that most studies target at DLPFC [14]. Stimulating the PPC have also been found improving the associative memory function [15–17]. However, tentative TMS targets for modulating the episodic memory function at cortical regions functionally connected to the hippocampus, to our knowledge, has not been explored. In this study, we aim at revealing the personalized TMS targets to preserve or even promote the episodic memory function. Particularly, we study the cortical regions that are functionally connected to the hippocampus in the resting state in a large cohort (Human Connectome Project [18]). Our goal is to reveal centroids and the spatial distribution of these cortical regions to inform 1) cortical areas that are TMS target candidates for modulating the episodic memory function and 2) coordinates to guide TMS when individual fMRI is not available. This information can explain the variability of the episodic memory function modulation by TMS and suggest methods to improve the modulation stability and effect.

## Materials and methods

The resting-state fMRI data for 1,133 young adults were downloaded from the Human Connectome Project database. The preprocessed data [19] had 2-mm isotropic spatial resolution and were sampled at the rate of 0.72 s per brain volume. Functional MRI data were spatially registered to the structural MRI data using FSL (*flirt)*. We used the seed-based correlation to delineate the functional connectivity to the hippocampus. Two approaches were taken to generate the seed time course: We averaged the fMRI time courses at the hippocampus automatically segmented from an individual’s structural MRI by FreeSurfer [20]. We denote this as the *Seed* approach in this study. Alternatively, the hippocampus fMRI time series was calculated by the *Seedmap* approach [21], which was a weighted spatial average of the fMRI time courses across all gray matter image voxels, excluding medial prefrontal and bilateral parietal regions. Weights were derived from the group-level functional connectivity map (see description below).

We used linear regression to remove temporal confounds in the fMRI time series related to the fluctuations in white matter and ventricles, which were automatically segmented by FreeSurfer using an individual’s structural MRI. The functional connectivity to the hippocampus was then created by calculating the Pearson correlation coefficients between the fMRI time courses at the hippocampus (either *Seed* or *Seedmap* approach) and at each brain location. The correlation coefficients were then transformed to Z scores using Fisher’s transformation [22]. In this calculation, we considered the temporal correlation of the fMRI time series based on Barlett’s theory [23], which estimated the number of independent measurements as *n*_t_ /9.8, where *n*_t_ denotes the number of samples in the fMRI time series. The spatial distribution of Z scores was taken as an individual’s functional connectivity map. The group-level functional connectivity was calculated by testing if the Z score was non-zero across all participants. Bonferroni correction was applied to suppress the inflation of the type-I error caused by multiple comparisons.

Three sets of TMS targets were created in this study. First, we used a large-scale database (http://neurosynth.org) to perform the automated meta-analysis. With the term “memory retrieval,” we identified targets at mPFC, left PPC, and right PPC at MNI coordinates of (0, 62, 4), (-42 -64, 48), and (40, -66, 44), respectively. These were taken as *atlas* targets in this study. Centroids of the group-level functional connectivity at the mPFC and bilateral PPC were taken as the group-level functional connectivity (*grp FC*) targets. The average centroids of each individual’s functional connectivity at the mPFC and bilateral PPC were taken as the individualized functional connectivity (*ind FC*) targets.

Given one set of TMS targets, we calculated the Euclidean distance between the centroid of an individual’s functional connectivity at mPFC and PPC to the TMS target. The bias and the dispersion of TMS targets were quantified by the average, standard deviation, median, and interquartile range of these distances across individuals.

With the calculated functional connectivity maps in each individual, we also calculated the penetration map to show the spatial distribution of the percentage of individuals showing significant functional connectivity to the hippocampus. Specifically, a binary mask was created for each individual to denote the areas of significant functional connectivity using either *Seed* or *Seedmap* approach after correcting the inflation of multiple comparisons by controlling the false discovery rate to 0.05. Averaging binary masks across individuals led to a sensitivity measure disclosing the proportion of a population showing significant functional connectivity to the hippocampus at each brain location.

## Results

Figure 1. shows maps of the group-level functional connectivity to the hippocampus. Using either *Seed* or *Seedmap* approach, we found functionally connected areas at the mPFC and bilateral PPC. In addition, the precuneus and bilateral temporal poles had correlated hemodynamics to the hippocampus. In the interest of studying TMS targets in brain regions close to the scalp, we only focus on the analysis of mPFC and PPC in this study.

**Figure 1.**
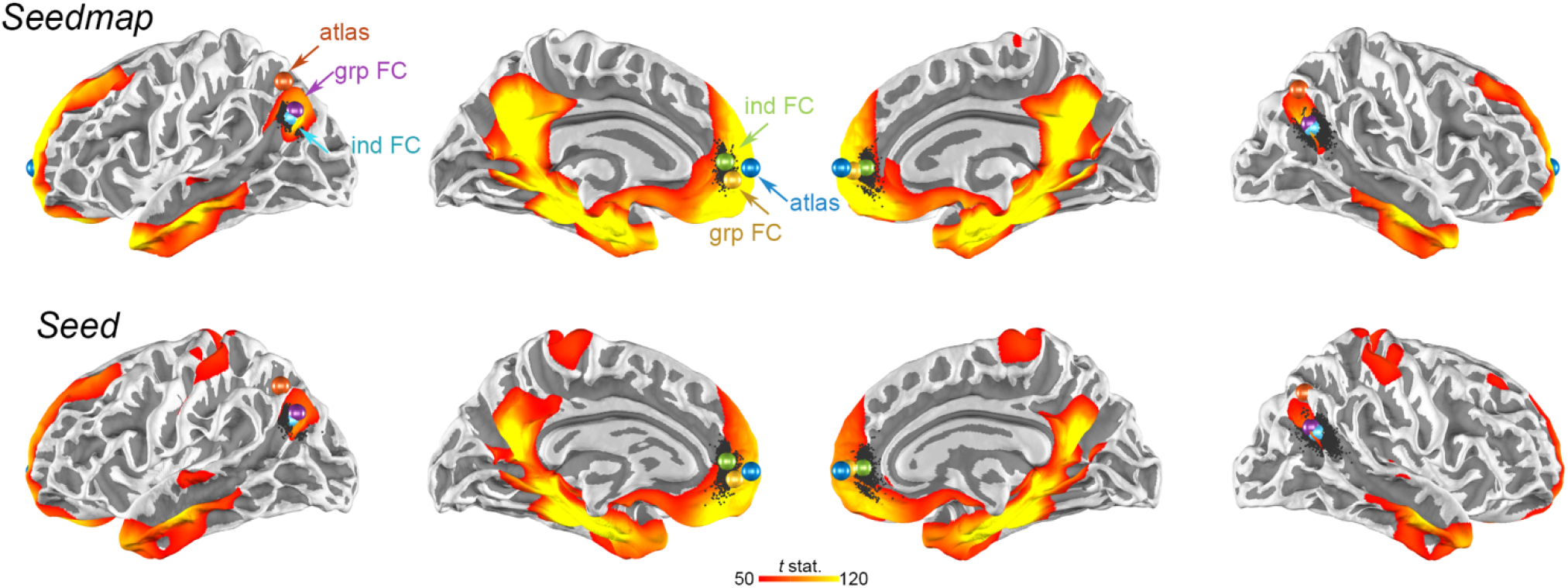
Maps of functional connectivity to the hippocampus using *Seedmap* (top) or *Seed* (bottom) to extract the hippocampal time series from a large cohort (*n* = 1,133). Color represents *t* statistics. Black dots are the centroids of the functional connectivity at the medial prefrontal cortex (mPFC) and posterior parietal cortex (PPC) in each individual. Centroids of the group-level functional connectivity (grp FC) at mPFC and PPC are represented by brown and purple spheres, respectively. Average of centroids of individual functional connectivity (ind FC) at mPFC and PPC are represented by green and cyan spheres, respectively. Atlas coordinates at mPFC and PPC are represented by blue and orange spheres, respectively.

Visually, centroids of the functional connectivity for each individual, shown as black dots in **Figure 1**, were more spatially dispersed at the mPFC than at PPC. In the left hemisphere, the group-level functional connectivity targets at mPFC and PPC were (-9, 54, 0) and (-41, -72, 34), respectively. In the right hemisphere, the group-level functional connectivity targets at mPFC and PPC were (10, 59, 5), and (47, -62, 26), respectively. The individualized functional connectivity targets at the left hemisphere were (-10, 50, 9) and (-39, -70, 30) for mPFC and PPC, respectively. These individualized targets at the right hemisphere were (11, 51, 6), and (48, -59, 24) for mPFC and PPC, respectively. In the left hemisphere, the distance between group-level targets and the atlas targets were 12.7 mm and 16.2 mm at mPFC and PPC, respectively. These distances were 10.5 mm and 19.7 mm in the right mPFC and right PPC, respectively. The distance between group-level targets and the atlas targets were 16.4 mm and 19.2 mm at mPFC and PPC at the left hemisphere, respectively. These separations were 15.7 mm and 22.6 mm at mPFC and PPC at the right hemisphere, respectively. Between group-level and individualized targets, the distances were 9.9 mm and 4.9 mm in the left hemisphere, respectively, and 8.1 mm and 3.7 mm in the right hemisphere, respectively. The locations of these targets are shown by colour spheres in **Figure 1**.

Targets estimated by *Seedmap* approach were similar to those estimated by *Seed* approach. Details of the target coordinates are reported in **Table 1**. Atlas targets were more than 10 mm away from functional connectivity targets from either group-level or individualized analysis at both mPFC and PPC. **Figure 2** shows distributions of the distances between TMS targets and individual’s functional connectivity centroids at mPFC and PPC. Briefly, the average of these distances were larger and their dispersions were less in functional connectivity-defined targets than atlas targets.The dispersion of the distance between an individual’s functional connectivity centroid and a TMS target, quantified by either the standard deviation or interquartile ratio of distances, at PPC and mPFC were in the range between 3 mm and 5 mm. This range suggested the probabilistic bounds for individualized TMS targets when no fMRI measurement is available. Between *Seed* and *Seedmap* approaches, the dispersion estimated by *Seedmap* has a smaller standard deviation and interquartile ratio, suggesting that *Seedmap* target coordinates are more stable across individuals.

**Table 1.** TMS target coordinates and measures (mean, median, standard deviation (Std), and interquartile range (IQR)) of the distance to TMS targets defined by atlas, group-level functional connectivity (grp FC), and individual functional connectivity (ind FC). Hippocampal time series defined by *Seed* and *Seedmap* approaches are reported separately.

|  |  | Target | mPFC |  |  |  |  |  |  | PPC |  |  |  |  |  |  |
| --- | --- | --- | --- | --- | --- | --- | --- | --- | --- | --- | --- | --- | --- | --- | --- | --- |
|  |  |  | MNI coordinate |  |  | distance to target (mm) |  |  |  | MNI coordinate |  |  | distance to target (mm) |  |  |  |
|  |  |  | X | Y | Z | mean | Std | median | IQR | X | Y | Z | mean | Std | median | IQR |
| Left hemisphere | Seed map | atlas | 0 | 62 | 4 | 18.1 | 1.5 | 17.9 | 1.8 | -42 | -64 | 48 | 18.9 | 4.2 | 18.2 | 5.8 |
|  |  | grp FC | -9 | 53 | -2 | 11.1 | 3.1 | 11.2 | 1.5 | -41 | -69 | 33 | 6.2 | 2.9 | 5.7 | 4.4 |
|  |  | ind FC | -10 | 49 | 7 | 3.4 | 2.3 | 2.8 | 2.6 | -40 | -67 | 30 | 4.8 | 2.6 | 4.4 | 3.6 |
|  | Seed | atlas | 0 | 62 | 4 | 18.2 | 2.0 | 18.1 | 2.1 | -42 | -64 | 48 | 19.2 | 5.1 | 18.5 | 7.6 |
|  |  | grp FC | -9 | 54 | 0 | 11.6 | 3.6 | 11.6 | 2.5 | -41 | -72 | 34 | 7.0 | 3.4 | 6.5 | 5.0 |
|  |  | ind FC | -10 | 50 | 9 | 4.5 | 3.3 | 3.7 | 3.7 | -39 | -70 | 30 | 5.5 | 3.0 | 5.0 | 4.1 |
| Right hemisphere | Seed map | atlas | 0 | 62 | 4 | 17.8 | 1.7 | 17.7 | 1.7 | 40 | -66 | 44 | 21.8 | 4.6 | 21.6 | 5.7 |
|  |  | grp FC | 10 | 56 | 2 | 8.2 | 2.4 | 7.9 | 2.3 | 45 | -60 | 27 | 5.8 | 2.9 | 5.1 | 4.0 |
|  |  | ind FC | 12 | 50 | 4 | 4.2 | 2.9 | 3.5 | 3.3 | 46 | -59 | 25 | 5.2 | 2.7 | 4.8 | 3.7 |
|  | Seed | atlas | 0 | 62 | 4 | 17.1 | 2.5 | 16.7 | 2.4 | 40 | -66 | 44 | 23.0 | 5.4 | 22.5 | 6.7 |
|  |  | grp FC | 10 | 59 | 5 | 9.9 | 3.4 | 9.1 | 3.1 | 47 | -62 | 26 | 7.0 | 3.8 | 6.0 | 5.1 |
|  |  | ind FC | 11 | 51 | 6 | 5.6 | 3.8 | 4.6 | 4.3 | 48 | -59 | 24 | 6.1 | 3.3 | 5.5 | 4.5 |

**Figure 2.**
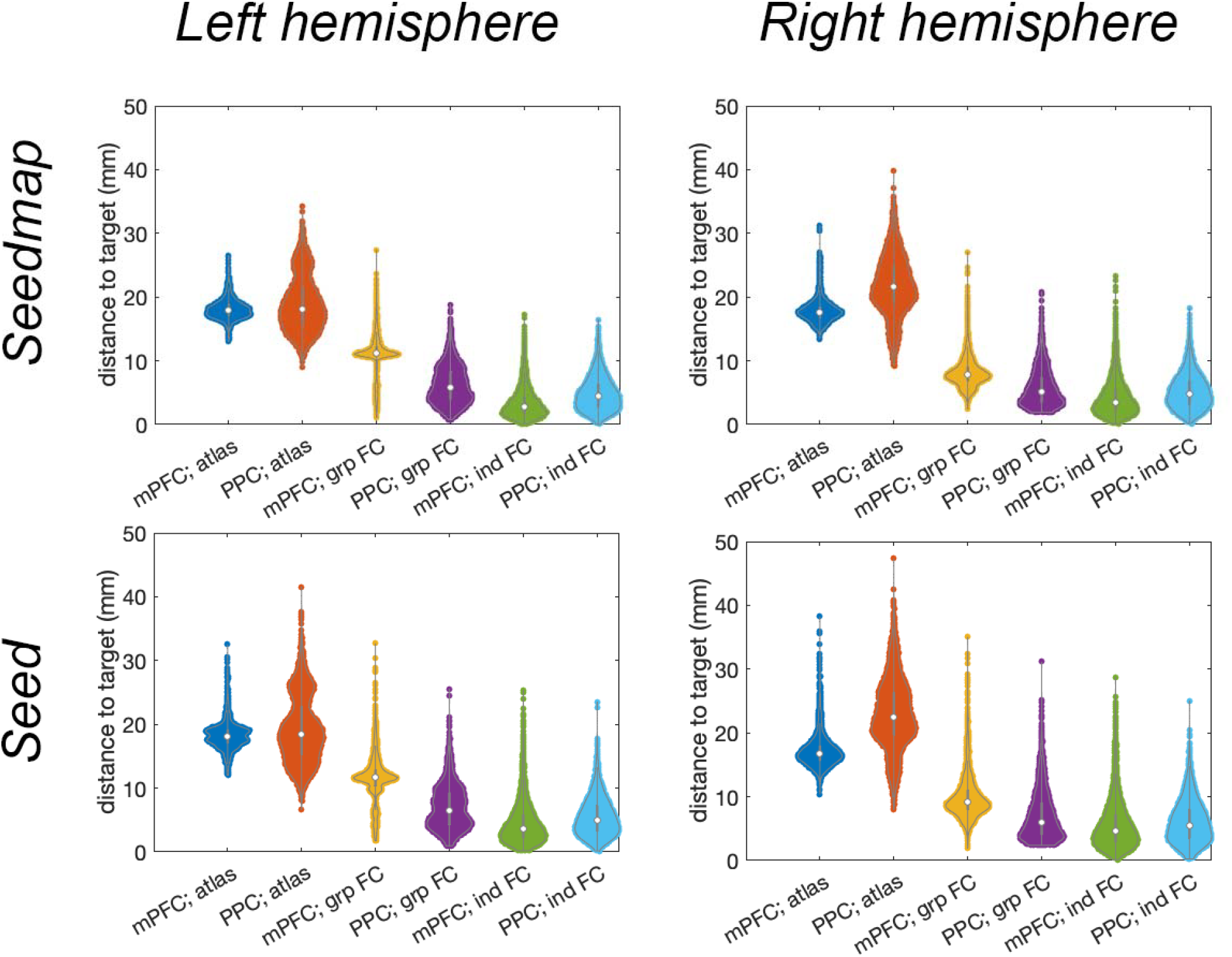
Violin plots of the distance between an individual’s functional connectivity centroid to TMS targets defined by group-level functional connectivity (grp FC), average of individual’s functional connectivity (ind FC), and atlas coordinates (atlas) at mPFC and PPC using *Seedmap* (top) and *Seed* (bottom) approach to describe the hippocampal time series.

Lastly, we examined the proportion of participants showing significant hippocampus functional connectivity maps using either *Seed* or *Seedmap* to generate the hippocampal time series. **Figure 3** shows these penetration maps. Only around 25% of participants showed functional connectivity to the hippocampus at mPFC and PPC with the *Seed* approach. In contrast, using *Seedmap* dramatically increases the proportion (>90%) of participants showing statistically significant mPFC and PPC regions connecting to the hippocampus, suggesting that *Seedmap* is more sensitive in revealing tentative TMS targets.

**Figure 3.**
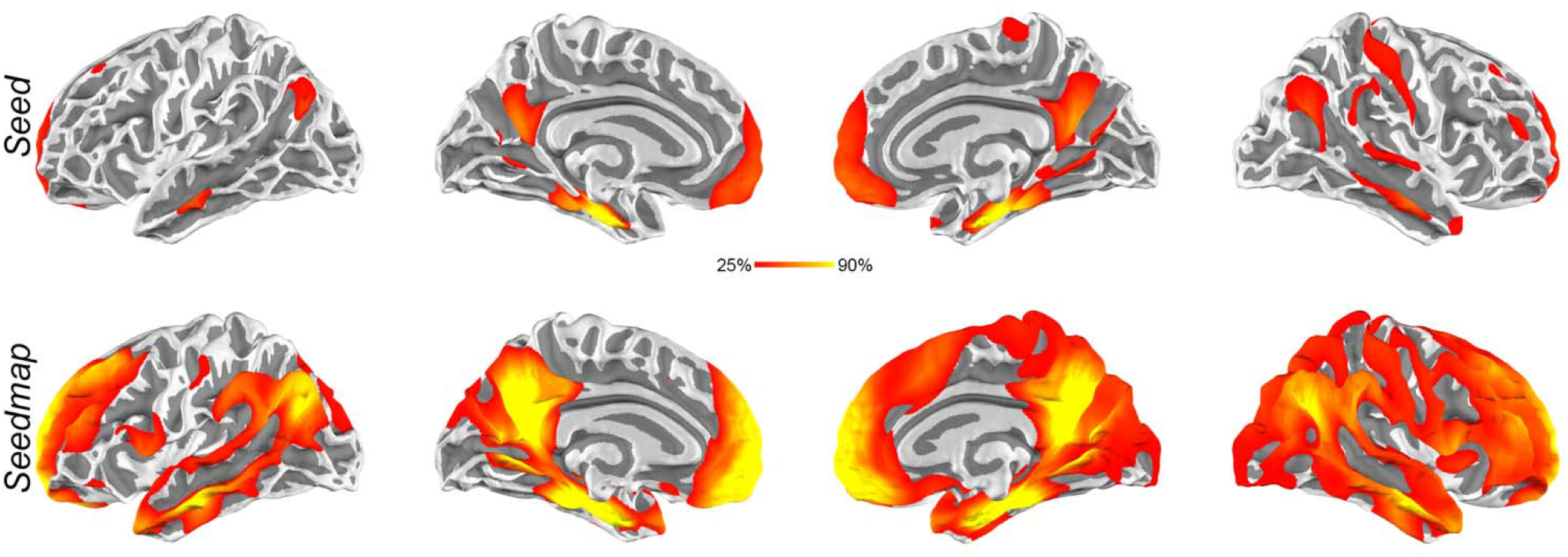
Penetration maps of functional connectivity using *Seed* (top) and *Seedmap* (bottom) approaches across 1,133 individuals. The color of each brain area represents the percentage of the population showing significant functional connectivity to the hippocampus.

## Discussion

Using fMRI data from a large cohort, we show that cortical areas functionally connected to the hippocampus varied across individuals. The present study was motivated by the recent view of using personalized TMS targets for the optimal modulatory effect [9]. Importantly, at either PPC or mPFC, a common TMS target can be away from the centroid of the functionally connected cortex for 10 mm or more. This discrepancy may be related to the remission of TMS due to the sub-optimal stimulation because the induced magnetic field strength drops to 50 % or less at 5 mm from the target [24].

The separation between functional connectivity-derived TMS targets and atlas TMS targets can be attributed to the difference in the brain activities between task engagement, which eventually led to atlas targets in the meta-analysis, and the resting state. Empirically targeting atlas and functional connectivity targets is required to validate which targets are more effective in preserving or promoting episodic memory function.

The difference in the sensitivity of revealing TMS targets at mPFC and PPC between *Seed* and *Seedmap* approaches (**Figure 3**) can be attributed to the fact that part of the brain dynamics within the hippocampus has been removed in *Seedmap* calculation, where only a weighted sum of cortical fMRI waveforms was used to represent the hippocampal dynamics. While *Seedmap* led to a more stable revelation of the functionally connected regions at mPFC and PPC across individuals, caution should be taken that such a “seed” time series was a surrogate of the hippocampal dynamics. Practically, within an individual, *Seedmap* is more likely to identify the TMS targets at mPFC and PPC than *Seed* with the same statistical threshold.

The functional connectivity-derived TMS targets are of practical value for studies without access to individual resting-state fMRI. The reported coordinates can suggest candidate TMS targets to modulate episodic memory function. In fact, these coordinates can also be effective in modulating other functions, such as learning and emotion, where the hippocampus plays a pivotal role.

Identifying mPFC and PPC coordinates functionally connected to the hippocampus corroborates a potential strategy of stimulating the hippocampus using multi-focus TMS [25,26]. Specifically, targeting both mPFC and PPC is likely to stimulate hippocampal activity. Separate TMS targeting at mPFC [27–29] and PPC [15,17,30] has already confirmed that hippocampal activity can be activated by non-invasive cortical stimulation. Applying TMS at two sites may synergically modulate hippocampal activity and memory function more effectively. Yet, the latency or phase delays between stimulations at mPFC and lPPC and the strength of the stimulation for optimal outcomes remain unknown.

The distribution of the functional connectivity locations (**Figure 1**) at mPFC and PPC can be useful in developing a new TMS coil, which can consider this spatial dispersion across individuals in depth-focality trade-off [8]. Together with the electric field maps, our calculation can also suggest the TMS coil orientations and locations, if no individual fMRI is available, to stimulate most participants’ hippocampal networks.

This study’s limitations include data from healthy young adults; thus, effects related to aging cannot be revealed. The data were only in the resting state, precluding potential functional connectivity variability in other conditions closely related to memory formation, consolidation, and retrieval. While our results suggest coordinates for TMS coil placement, we cannot suggest the TMS coil orientation, an important parameter to induce the optimal electric field [31]. This gap can be bridged by electric field modeling with realistic brain anatomy [24] constrained by our suggested coordinates.

